# Intraepithelial T cells diverge by intestinal location as pigs age

**DOI:** 10.1101/2020.03.05.973248

**Authors:** Jayne E. Wiarda, Julian M. Trachsel, Zahra F. Bond, Kristen A. Byrne, Nicholas K. Gabler, Crystal L. Loving

## Abstract

T cells resident within the intestinal epithelium play a central role in barrier integrity and provide a first line of immune defense. Intraepithelial T cells (IETs) are among the earliest immune cells to populate and protect intestinal tissues, thereby giving them an important role in shaping gut health early in life. In pigs, IETs are poorly defined, and their maturation in young pigs has not been well studied. Given the importance of IETs in contributing to early life and long-term intestinal health through interactions with epithelial cells, the microbiota, and additional environmental factors, a deeper characterization of IETs in pigs is warranted. The objective of this study was to analyze age- and intestinal location-dependent changes in IETs across multiple sites of the small and large intestine in pigs between 4 and 8 weeks of age. IETs increased in abundance over time and belonged to both γδ and αβ T cell lineages. Similar compositions of IETs were identified across intestinal sites in 4-week-old pigs, but compositions diverged between intestinal sites as pigs aged. CD2^+^CD8α^+^ γδ T cells and CD4^−^CD8α^+^ αβ T cells comprised >78% of total IETs at all intestinal locations and ages examined. Greater percentages of γδ IETs were present in large intestine compared to small intestine in older pigs. Small intestinal tissues had greater percentages of CD2^+^CD8α^−^ γδ IETs, while CD2^+^CD8α^+^ γδ IET percentages were greater in the large intestine. Percentages of CD4^−^CD8α^+^ αβ IETs increased over time across all intestinal sites. Moreover, percentages of CD27^+^ cells decreased in ileum and large intestine over time, indicating increased IET activation as pigs aged. Percentages of CD27^+^ cells were also higher in small intestine compared to large intestine at later timepoints. Results herein emphasize 4 to 8 weeks of age as a critical window of IET maturation and suggest strong associations between intestinal location and age with IET heterogeneity in pigs.

## 1 Introduction

The intestinal tract contains the single largest and most diverse compartment of immune cells in the body and is a highly versatile organ system, with different regions performing various physiological and immune functions (1). Furthermore, the intestinal epithelium performs a key function by serving as a selective physical barrier between the intestinal lumen and the body. Intraepithelial T cells (IETs) are T cells located within the epithelial layer throughout the intestinal tract. In neonates, IETs are among the earliest immune cells to populate and protect intestinal tissues (2). IETs are tissue-resident cells that respond to foreign antigen from the intestinal lumen and to self-derived, stress-induced molecules, positioning IETs as first responders to enteric pathogens or as mediators of epithelial stress, respectively (2-5). IETs are also important in regulating metabolism; mice lacking IETs are resistant to weight gain, even when fed a high-fat diet, due to a hyperactive metabolic profile, indicating IETs have an important role in promoting weight gain efficiency (6). In the context of immunity, IETs can release cytotoxic molecules, cytokines, and/or antimicrobial peptides upon activation, inducing intestinal inflammation and/or antimicrobial activity associated with immune protection (7-9). Conversely, to avoid unwarranted inflammation, IETs are tightly regulated and exercise regulatory functions, giving them a primary role in maintaining epithelial integrity and immune quiescence (7, 8). In the absence of such regulation, inflammation induced or exacerbated by IETs can threaten epithelial barrier integrity and promote immunopathology (2). Hence, IETs are critical in balancing immune tolerance and protection within the intestinal tract. Moreover, microbial and dietary antigen exposure majorly influence the development, specificity, reactivity, and homeostasis of IETs (6, 10-14), largely contributing to their fate in promoting or deteriorating intestinal health (2).

In humans and rodents, IET prevalence, phenotype, and function varies by anatomical location along the intestinal tract, indicating regional specialization of intestinal IETs, especially between small and large intestinal locations (15-20). Additionally, age is a primary driver of changes to intestinal IETs, indicating time-dependent changes to the cells occur due to further immune maturation and continual antigen exposure (15, 19, 21, 22). In pigs, information pertaining to IETs across intestinal locations or in regards to the impact of age during intestinal immune maturation is limited. While multiple studies have analyzed porcine T cells across different intestinal locations or across ages (23-29), the focus was primarily the small intestine and included combined cell fractions from epithelial and lamina propria compartments. Given the specialized role of intestinal IETs, it is important to understand whether previous findings can be attributed to intraepithelial versus lamina propria T cells or generalized to additional intestinal locations. In general, we know porcine small intestinal IETs are located primarily within the apical and middle portions of the villi, and the number of IETs increases with age, primarily in the first 3 months post-parturition, in a microbiota-dependent fashion (30-33). Similar to humans and rodents, the majority of IETs in the porcine small intestine are CD4^−^ CD8α^+^ (34); however, whether IETs belong to αβ or γδ T cell lineages is unknown and could have further implications into cell function. Meanwhile, studies analyzing IETs within the porcine large intestine are lacking.

In pigs, the window of approximately 3 to 8 weeks of age (often referred to as the weaning and nursery period in pig production) is a critical time during which porcine intestinal T cell communities are still developing (25, 35-37). Stress from the weaning process, as pigs are moved from the dam and a milk-based diet to new surroundings, pen mates, social structure, and solid food, can result in intestinal inflammation, increased epithelial permeability, diarrhea, increased susceptibility to disease, decreased nutritional absorption, and weight loss with life-long effects (38, 39). A better understanding of age- and location-dependent characteristics of intestinal IETs during stages of major immune maturation and increased stress, such as that of the nursery phase, may prove useful in developing strategies to improve pig health and/or market performance. To our knowledge, an analysis of age- and intestinal location-dependent changes in porcine αβ and γδ IET abundance, phenotype, and distribution throughout multiple compartments of both small and large intestine during intestinal T cell maturation in nursery-age pigs has not been completed. Hence, the objective of this study was to quantify IET numbers, assess presence and proportional phenotypes of both αβ and γδ IET populations, and assess expression of the T cell activation marker CD27 between jejunal, ileal, cecal, and colonic tissues across multiple weeks of age in pigs during the nursery period.

## 2 Materials and Methods

### 2.1 Study overview

Conventional, mixed-breed pigs were weaned from dams at ∼19-21 days of age and transported to the Iowa State University Swine Nutrition Facility. Upon arrival, pigs were randomly selected, weighed, and individually penned. All pigs had free access to water and feed at all times. Pigs were fed a corn-soybean meal-based diet that met or exceeded nutrient and energy requirements for this size pig (NRC, 2012 #146). The diet was free of antibiotics and therapeutic concentrations of minerals. At ∼4, 6, and 8 weeks of age (7, 21, and 35 days post-weaning, respectively), pigs were randomly chosen, humanely euthanized, and exsanguinated. Immediately thereafter, intestinal tissue samples were collected. The study was completed in 2 identical replicates, with 4 pigs necropsied at each timepoint per replicate (n = 8 pigs per timepoint; n = 24 total).

### 2.2 Sample collection

Sections of jejunum, ileum, cecum, and colon were collected for tissue fixation and flow cytometric (FCM) staining. Jejunal sections were collected ∼95 cm distal to the pylorus. The most proximal ∼7.5-cm jejunal section was collected for tissue fixation, and the next ∼7.5-cm jejunal section was collected for FCM staining. Ileal sections were collected starting ∼7.5 cm proximal to the ileocecal valve. The more distal ∼7.5 cm ileal section was used for FCM staining, and the next ∼7.5 cm ileal section was collected for tissue fixation. Cecal sections were collected as two adjacent ∼5-cm by ∼10-cm sections located in the middle of the cecal pouch, one section for FCM staining and one section for tissue fixation. Colonic sections were collected from the apex of the spiral colon as two adjacent ∼7.5-cm colonic sections for FCM staining and tissue fixation.

### 2.3 Immunohistochemistry (IHC)

Intestinal tissues were fixed in a 10% neutral-buffered formalin solution (3.7% formaldehyde) for ∼24 hours at room temperature (RT). Tissues were then cut to appropriate size, placed in cassettes, transferred to 70% ethanol, and embedded in paraffin blocks. Formalin-fixed, paraffin-embedded (FFPE) tissues were cut into 4-micron thick sections and adhered to Superfrost-Plus charged microscope slides (Thermo Fisher Scientific). Immunohistochemical staining was performed for detection of CD3 protein as described previously (40). Briefly, slides were baked, deparaffinized, and rehydrated for IHC staining. Antigen retrieval was carried out by incubating slides in 1X sodium citrate buffer, pH 6.0 at 95 °C for 20 minutes in a pressurized Decloaking Chamber NxGen (Biocare Medical, LLC) and then allowing slides to cool down in antigen retrieval solution for ∼10 minutes outside of the decloaking chamber. Next, slides were sequentially incubated with endogenous enzyme blocker (Dako S2003) for 10 minutes at RT; protein block (Dako X0909) for 20 minutes at RT; 0.006 g/L polyclonal rabbit anti-human CD3 antibody (Dako A0452, stock concentration 0.60 g/L diluted 1:100 in 1% bovine serum albumin [BSA] phosphate-buffered saline [PBS]) for 60 minutes at RT; horseradish peroxidase (HRP)-labelled anti-rabbit antibody (Dako K4003) for 30 minutes at RT; and 3,3’-diaminobenzidine (DAB) substrate (Dako K3468) for 3 minutes at RT. Volumes used for each incubation varied between slides but was enough to fully cover all tissue sections. Between each incubation, slides were washed with 0.05% PBS-Tween (PBS-T), pH 7.35±0.02. Slides were then counterstained with Gill’s Hematoxylin I (American Mastertech) for 1 minute, rinsed with distilled water, dehydrated, and coverslipped.

### 2.4 Dual Chromogenic IHC and RNA in-situ hybridization (IHC/ISH)

Dual chromogenic IHC/ISH staining was performed to simultaneously detect T receptor delta constant (*TRDC*) mRNA and CD3 protein. FFPE intestinal tissues were fixed, processed, and sectioned as described in IHC methods. Slides were first stained for *TRDC* mRNA using the RNAscope 2.5 HD Reagent Kit-RED (Advanced Cell Diagnostics, ACD) and custom-designed probe complementary to *Sus scrofa TRDC* mRNA (ACD 553141). A probe targeting *Bacillus subtilis DAPB* (ACD 310043) was used as a negative control. Slides were baked at 60 °C in a dry oven for 1 hour, followed by deparaffinization and rehydration using incubations in xylenes (2 × 5 minutes), 100% ethanol (2 × 1 minute), and air drying at RT. Slides were incubated with Hydrogen Peroxide (ACD) for 10 minutes at RT, rinsed with water, incubated in 1X Target Retrieval Solution (ACD) for 15 min at 95 °C in a pressurized Decloaking Chamber NxGen (Biocare), rinsed with distilled water, incubated in 100% ethanol for 2 minutes, and air dried at RT. Once dry, a hydrophobic barrier was drawn around each tissue using an ImmEdge PAP pen (Vector Laboratories, Inc.).

ISH staining for *TRDC* RNA was completed by incubating slides in a humidifying tray either at at 40 °C in a HybEZ Hybridization System oven (ACD) or at RT on the benchtop for all steps. Protein digestion was performed by incubating slides with Protease Plus (ACD) for 15 minutes at 40 °C, followed by rinsing with distilled water. Next, slides were sequentially incubated with the following reagents and washed with 1X Wash Buffer (ACD) 2 × 2 minutes between each incubation: undiluted *TRDC* probe (ACD) 2 hours at 40 °C; 5X saline-sodium citrate (SSC) buffer overnight at RT; AMP1 (ACD) at 40 °C for 30 minutes; AMP2 (ACD) at 40 °C for 15 minutes; AMP3 (ACD) at 40 °C for 30 minutes; AMP4 (ACD) at 40 °C for 15 minutes; AMP5 (ACD) at RT for 30 minutes; AMP6 (ACD) at RT for 15 minutes; and prepared RED detection solution (diluted according to manufacturer’s instructions; ACD) at RT for 10 minutes.

Following RNA ISH, IHC was performed for CD3 protein staining. Slides were washed with 0.05% PBS-T, pH 7.35±0.02 (2 × 2 minutes) following RNA ISH and following incubations with protein block, primary antibody, and secondary antibody as outlined in the CD3 IHC method. Next, slides were incubated with HIGHDEF Yellow HRP chromogen (diluted according to manufacturer’s instructions; Enzo Life Sciences) for 10 minutes at RT followed by washing again with PBS-T. To counterstain, slides were placed into 25% Gill’s Hematoxylin I (American Mastertech) for 30 seconds. Following counterstaining, slides were rinsed well with distilled water, dried for 20 minutes at 60 °C, and mounted with VectaMount Permanent Mounting Media (Vector) and #1 thickness coverslips.

### 2.5 IHC stain quantification

Quantification of CD3 IHC staining within the intestinal epithelium of tissues was performed using the HALO image analysis platform (Indica Labs). Regions of interest were manually annotated around epithelium of 3 villi of jejunal and ileal tissues and 3 crypts of cecal and colonic tissues per sample. Only crypts or villi non-adjacent to mucosal-associated lymphoid tissue (e.g. Peyer’s patches) were annotated for analysis. Due to the inability to accurately define individual cell borders of tightly-packed cells using the software, CD3 staining was quantified as a percentage of CD3-stained surface area over the total surface area of the annotated regions with user-defined parameters for stain detection from the Area Quantification (v2.1.3) package. The percentage of CD3-stained surface area for each of the 3 annotated villi or crypts per sample were averaged together to obtain a single value for each sample. Software quantification of dually-stained CD3 and *TRDC* in tissues was not performed due to cross-detection between the two chromogenic stains.

### 2.6 Analysis of variance (ANOVA) statistical analyses of IHC data

One-way ANOVA analyses of percentages obtained from IHC staining quantification were performed using Prism 8 (version 8.1.2; GraphPad Software). A Gaussian distribution could not be assumed based on small sample size; therefore, nonparametric tests were used to analyze data. The rank-based Kruskal-Wallis test was performed on sets of data within a single tissue across timepoints. All combinations of multiple comparisons between tissues or timepoints were analyzed within each dataset for both analyses. P-values < 0.05 were considered significant (* < 0.05, ** < 0.01, *** < 0.001, **** < 0.0001). Input data can be found in supplemental materials.

### 2.7 Intestinal epithelial isolation

Sections of jejunum, ileum, cecum, and colon were collected to obtain single-cell suspensions for cell phenotype labeling and analysis by FCM. Intestinal sections were cut open to expose the lumen, and the epithelium was gently rinsed with PBS to remove intestinal contents. Sections were placed in RT stabilization buffer of Hank’s balanced salt solution (HBSS; Gibco 14175) containing 2 mM ethylenediaminetetraacetic acid (EDTA; Invitrogen AM9261), 2 mM L-glutamine (Gibco 25030), and 0.5% BSA (Sigma A9418) for transport back to the lab. In the lab, ∼1.5-gram sections of tissues were processed for single-cell isolates, and all subsequent incubations were performed in a shaking incubator (200 rpm, 37 °C). Mucus dissociation was performed by incubating tissues in 30 mL of HBSS containing 5mM dithiothretol (DTT; Invitrogen 15508) and 2% heat-inactivated fetal calf serum (FCS; Gibco A38401) for 20 minutes. Epithelial cell removal was carried out by transferring tissue into 30 mL of HBSS containing 5 mM EDTA and 2% FCS. A total of 3 sequential incubations in fresh epithelial removal solution were carried out for 25 minutes each, transferring the tissue to fresh solution for each incubation. Tissues were then washed in 20 mL of HBSS containing 10 mM HEPES (Fisher Scientific BP299) for 10 minutes before transferring to 10% neutral buffered formalin to confirm epithelial cell removal. Liberated cells from the epithelial removal and wash solutions were retained, pooled, passed through a 100-micron nylon filter, and washed with HBSS containing 2 mM L-glutamine and 2% FCS. Isolated cells were centrifuged 8 minutes at 450 × g at RT, and the pellet was resuspended in HBSS/L-glutamine/FCS solution. Viability and quantity of the final epithelial-enriched cell suspensions were determined with the Muse Cell Analyzer with the Muse Count & Viability Assay Kit (Luminex).

### 2.8 Cell phenotype labeling and data acquisition by flow cytometry

For each sample, 5×10^5^ live cells were seeded into a single well of a 96-well round bottom plate, pelleted, resuspended, and stained with Fixable Viability Dye eFluor 780 (eBioscience) diluted 1:1000 in PBS for 30 minutes according to manufacturer’s recommendations. Next, sequential incubations with unconjugated primary antibodies, secondary fluorophore-conjugated antibodies, and primary antibodies directly-conjugated to fluorophores were carried out for 15 minutes each at RT. Unconjugated primary antibodies included anti-γδ T cell receptor (α-γδTCR; PGBL22A, mouse IgG_1_) and α-CD2 (MSA4, mouse IgG_2a_) from Washington State University. Secondary antibodies included rat α-mouse IgG_1_-BUV395 (A85-1; BD Biosciences) and rat α-mouse IgG_2a_-BV605 (R19-15; BD). Directly conjugated antibodies included α-CD3ε-PE-Cy7 (BB23-8E6-8C8, mouse IgG_2a_; BD), α-CD4-PerCP-Cy5.5 (74-12-4, mouse IgG_2b_; BD), α-CD8α-PE (76-2-11, mouse IgG_2a_; BD), and α-CD27-FITC (b30c7, mouse IgG_1_; BioRad). Between incubations, cells were washed with PBS. After antibody staining, cells were fixed with BD Stabilizing Fixative (BD) and stored at 4 °C overnight. The following day, fixed cells were resuspended, passed through a 35-micron nylon filter to remove aggregates, and data were acquired using a BD FACSymphony A5 flow cytometer (BD). The instrument was set up according to manufacturer’s recommendations using bead capture reagents to set compensation controls.

### 2.9 Flow cytometry gating analysis

FCM data were analyzed with FlowJo (FlowJo, LLC). Single stains and fluorescence-minus-one antibody combinations were used to set appropriate gates for each fluorochrome (41, 42). Total IET communities of interest were defined as cells found within a lymphocyte size- and complexity-specific gate using SSC-A and FSC-A value plotting; singlets based on approximate 1:1 plotting of FSC-A versus FSC-H values; live cells based on negative fluorescence values for cell viability dye staining; and T cells based on positive fluorescence values for CD3ε-specific staining. Total IETs were further categorized as γδ or αβ IETs by having positive or negative fluorescence values for γδTCR-specific staining, respectively. γδ IETs were categorized into 4 gating quadrants based on positive or negative fluorescence values for CD2- and CD8α-specific staining. αβ IETs were categorized into 4 gating quadrants based on positive or negative fluorescence values for CD4- and CD8α-specific staining. The combined 4 γδ and 4 αβ IET quadrants gave rise to 8 IET populations. Within the 8 IET populations, CD27-specific fluorescence was also assessed to classify cells as CD27^+^ or CD27^−^, giving rise to a total of 16 IET subpopulations (8 IET populations × 2 CD27 classifications) at the highest resolution analyzed from our gating strategy. Data were quantified as frequencies of specified parent populations or total cell counts. Samples with low event yields were analyzed for outlier data using box-and-whisker plot outlier analysis for frequency measurements collected. If no outliers were detected, samples were included in further analysis.

### 2.10 t-distributed stochastic neighbor embedding (t-SNE) visualization of flow cytometry data

Dimensionality reduction using t-SNE visualization was performed on FCM data within FlowJo using the t-SNE and DownSample plug-ins available from FlowJo Exchange (https://www.flowjo.com/exchange/#/). Prior to t-SNE visualization, similar fluorescence intensities and gating for flow cytometry markers in each tissue at each timepoint were confirmed. Cells from the CD3ε^+^ gate of each sample were down-sampled (n = 990 cells per sample) using the DownSample plug-in to obtain equal numbers of cells for each sample type (based on combination of 4 tissues and 3 timepoints; 12 sample types total), and subsequent gates were reapplied to down samples. Next, the 8 gated IET populations were concatenated between all down samples within each tissue/timepoint combination to create a total of 96 concatenated files (8 files belonging to each IET population × 12 belonging to each tissue/timepoint combination). Keyword value series were applied based on IET population and sample type. Compensation was reapplied to the concatenated files, and the 96 files were again concatenated into a single file, including the keyword value series as additional parameters. Compensation was reapplied again to the final concatenation, and t-SNE analysis was completed with input parameters for fluorescence intensities of compensated CD3ε, γδTCR, CD2, CD4, CD8α, and CD27 being considered for visualization. Default options for the opt-SNE learning configuration, exact KNN algorithm, and Barnes-Hut gradient algorithm were used with 1,000 iterations and a perplexity of 150. To identify αβ and γδ IET populations within the final concatenation, gates were drawn based on the keyword series values assigned to IET populations and tissue/timepoint variables. CD27^+^ and CD27^−^ expression was assessed by redrawing gates used previously.

### 2.11 Non-metric multidimensional scaling (NMDS) visualization and permutational multivariate analysis of variance (PERMANOVA) statistical analyses of flow cytometry data

For a multivariate comparison of sample similarity, the IET compositions of each sample were considered. IET communities were composed of 16 discrete subpopulations defined by expression of CD27 (positive or negative) for each of the 8 T cell populations defined by flow cytometry gating analysis. Frequencies of the 16 subpopulations were calculated from cell counts exported from FlowJo and were considered as discrete, non-overlapping groups comprising the total CD3ε^+^ IET community within each sample. From the frequency data, a dissimilarity matrix was calculated using the Bray-Curtis dissimilarity metric and this dissimilarity matrix was used for visualization and statistical testing of sample similarity. NMDS visualization of Bray-Curtis dissimilarities was performed in R with the vegan (version 2.5-5) (43), and tidyverse (version 1.2.1) (44) packages. PERMANOVA testing was completed in R using vegan’s ‘adonis’ function. Tissue, age, and a combination of tissue and age were used as variables. Post-hoc pairwise PERMANOVA tests were completed on comparisons of interest (single tissue type between timepoints or within a single timepoint between tissues), correcting p-values with the false discovery rate (FDR) method. Corrected p-values < 0.05 were considered significant (* < 0.05, ** < 0.01, *** < 0.001, **** < 0.0001). Input data and R scripts can be found at https://github.com/jwiarda/Intraepithelial_T_cells.

### 2.12 ANOVA statistical analyses of flow cytometry data

One-way ANOVA using percentages obtained from individual gates of flow cytometry data were performed using Prism 8. Because a Gaussian distribution could not be assumed based on small sample size, nonparametric tests were used to analyze data. The rank-based Kruskal-Wallis test was performed on sets of data within a single tissue across timepoints, while the paired, rank-based Friedman test was performed on sets of data within a single timepoint across tissues, pairing samples derived from the same animal. All combinations of multiple comparisons between tissues or timepoints were analyzed within each data set for both analyses. P-values < 0.05 were considered significant (* < 0.05, ** < 0.01, *** < 0.001, **** < 0.0001). Input data can be found in supplemental materials.

## 3 Results

### 3.1 Intestinal IET abundance increased with age and was composed of both γδ and αβ T cells

Jejunum, ileum, cecum, and colon were collected from 4-, 6-, and 8-week-old pigs (Fig. 1A). To assess the presence of T cells in the intestine, IHC staining of CD3 protein was completed using the collected tissues, and any CD3 stain-positive cells were considered T cells. T cells were found in both the epithelial layer and the lamina propria of all intestinal tissues, as well as within the Peyer’s patch areas of the ileum. CD3 staining within the epithelium appeared to be more frequent in the villi compared to the crypts of small intestinal tissues (jejunum and ileum) and within the apical portions of crypts in large intestinal tissues (cecum and colon). Moreover, T cell staining within the epithelium appeared more frequent overall in small intestinal compared to large intestinal tissues and within a respective intestinal tissue as animals aged (Fig. 1B). To confirm the latter observation, CD3 staining was quantified within villus epithelium of jejunal and ileal tissues or crypt epithelium of cecal and colonic tissues and compared across time (Supplemental Fig. 1 & Supplemental Table 1). CD3 staining within the epithelium increased significantly at all intestinal sites as age increased (Fig. 1C), suggesting IETs became more abundant across the 4 to 8 weeks of age time frame.

**Figure 1.**
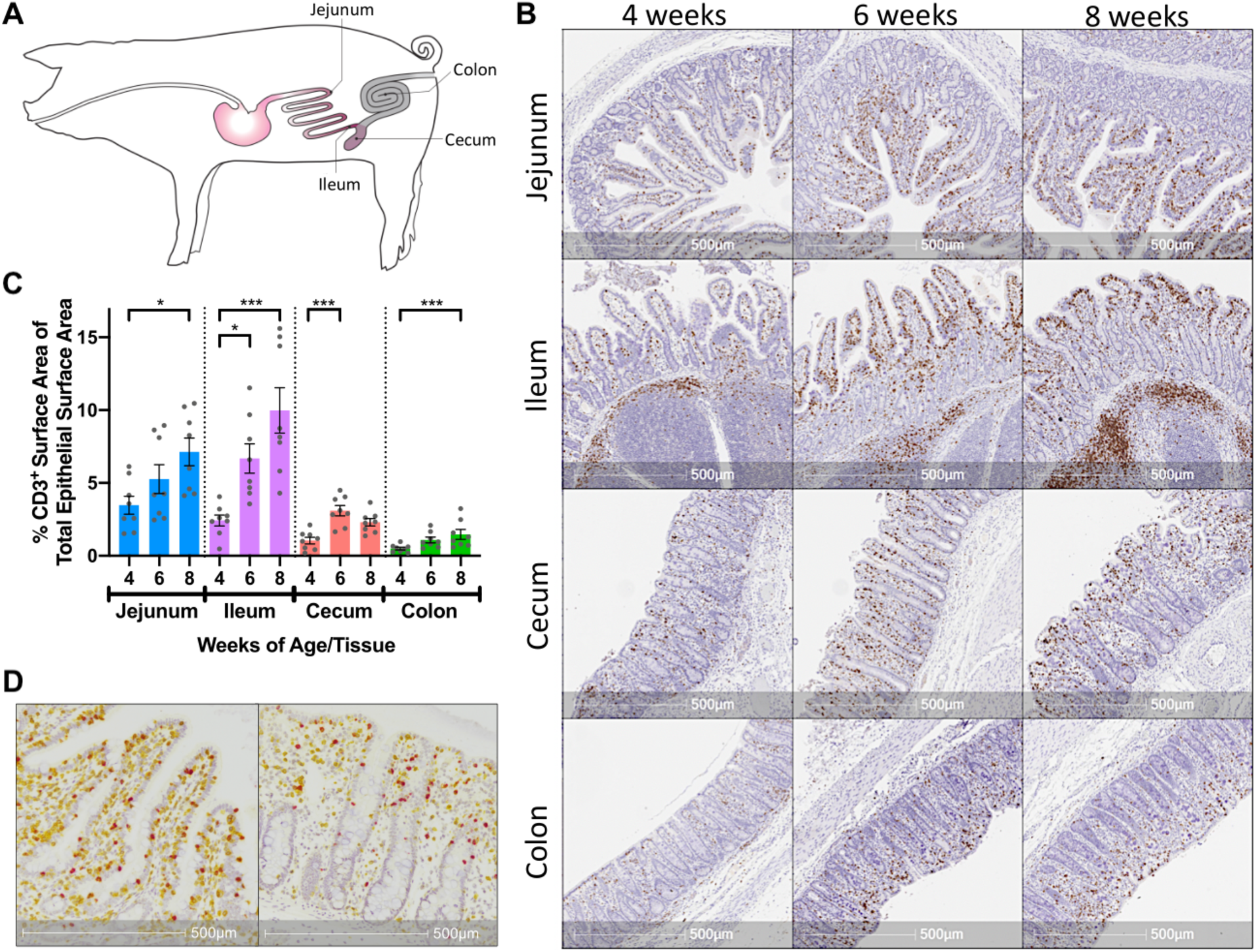
Porcine intestinal IETs increased with age and were composed of γδ and αβ T cells. **(A)** Anatomical diagram of tissue collection sites from jejunum, ileum, cecum, and colon within pig intestinal tract. **(B)** Representative images of T cell-specific CD3 protein staining (brown) within multiple intestinal tissues at different ages in pigs. **(C)** Percentages of epithelial surface area stained for CD3 protein within a single tissue over multiple timepoints. Statistical significance was determined within a single tissue across timepoints by the rank-based Kruskal-Wallis test using all combinations of multiple comparisons within a tissue. P-values < 0.05 were considered significant (* < 0.05, ** < 0.01, *** < 0.001, **** < 0.0001). **(D)** Representative images of T cell-specific CD3 protein staining (yellow) and γδ T cell-specific *TRDC* transcript staining (red) in pig ileum (left) and cecum (right) at 8 weeks of age. Distance scales on the bottom of each image are included for size comparisons. Samples were taken from 4 animals per timepoint in each of 2 separate experiments (n = 8 per timepoint; n = 24 total).

To further characterize intestinal IETs, dual staining of CD3 protein and T receptor delta constant (*TRDC*) mRNA was completed in a subset of jejunal, ileal, cecal, and colonic tissues. Using the rationale that CD3 protein (yellow staining) would be expressed by all T cells, *TRDC* mRNA (red staining) would only be expressed by γδ T cells, and that *TRDC*-specific red staining would mask co-localizing CD3-specific yellow staining, cells staining red were presumably γδ T cells (*TRDC*^+^), whereas cells staining yellow were presumably αβ T cells (CD3^+^*TRDC*^−^). Staining revealed the presence of both *TRDC*^+^ and CD3^+^*TRDC*^−^ cells, corresponding to presumable γδ and αβ T cells, respectively, within the epithelium of all intestinal tissues analyzed (Fig. 1D). T cells in the lamina propria and submucosa were primarily CD3^+^*TRDC*^−^, with *TRDC*^+^ cells noted infrequently outside of the epithelium (Supplemental Fig. 2). These findings suggest both αβ and γδ T cells are located within the intestinal epithelium of both small and large intestine in pigs.

### 3.2 IET compositions diverged by intestinal location as pigs aged

To further phenotype intestinal IETs, cell staining and FCM analysis was performed on epithelial-enriched fractions of the jejunum, ileum, cecum, and colon collected at 4, 6, and 8 weeks of age. The protocol used to liberate epithelial cells from the intestinal tissues resulted in isolation of primarily epithelial cells, though some lamina propria cells may have been released during processing and included in the analysis (Supplemental Fig. 3). CD3ε was used to identify T cells, and further characterization of total T cells by expression of cell surface markers γδTCR, CD2, CD4, CD8α, and CD27 was assessed. CD2 and CD8α expression are commonly used to identify functional porcine γδ T cell populations (45, 46); CD4 and CD8α expression identify functional porcine αβ T cell populations (47, 48); and CD27 expression is indicative of T cell activation and memory states (49-52). In total, this gating strategy yielded 16 discrete, non-overlapping IET subpopulations based on CD27^+/-^ expression within each of the 8 defined IET populations (Fig. 2A).

**Figure 2.**
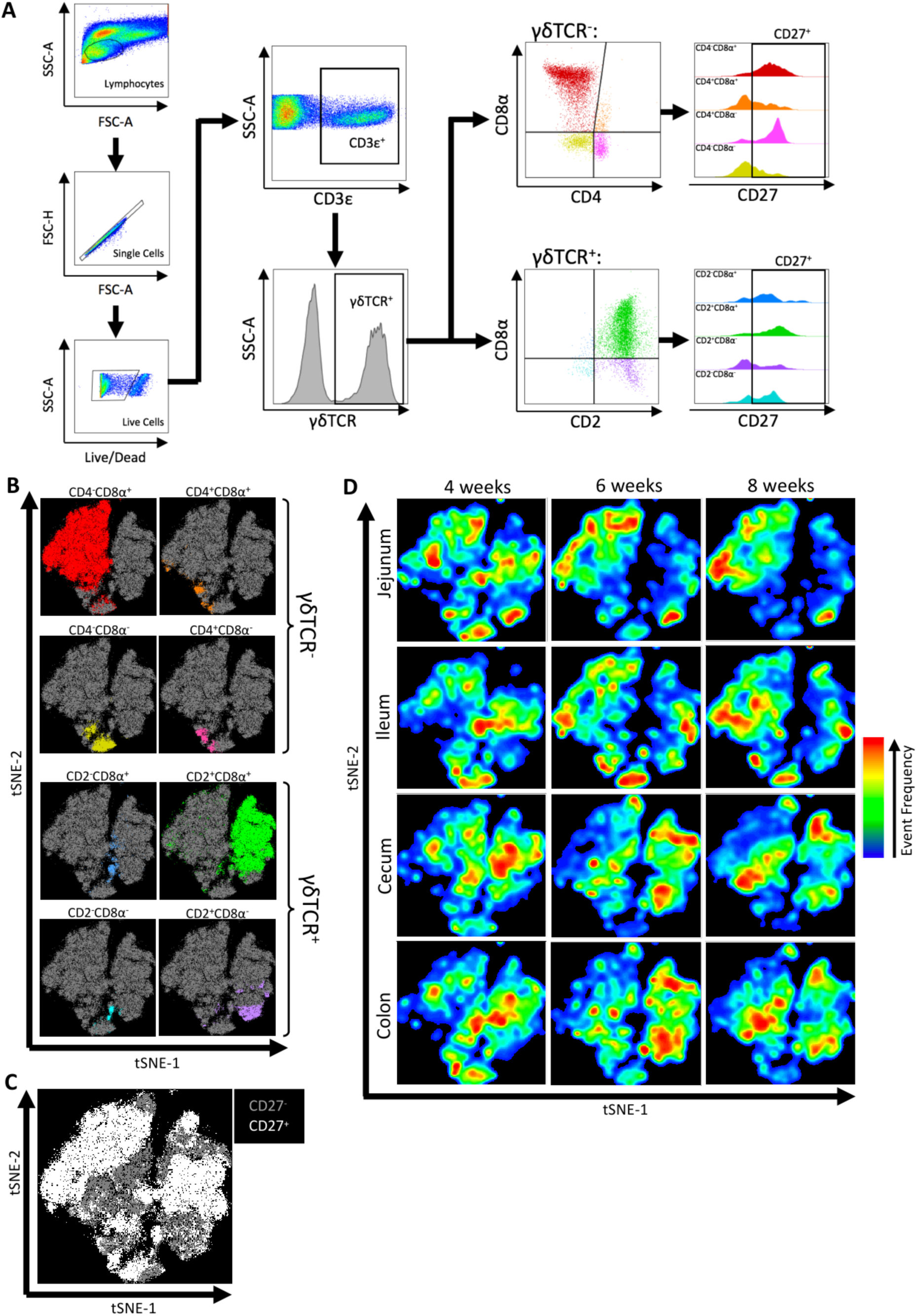
Flow cytometry gating revealed IET populations and CD27 expression varied by intestinal location and age. **(A)** Gating strategy used to identify IET populations and subpopulations from flow cytometry data of epithelial-enriched samples. IETs were identified by SSC-A and FSC-A profiles, single cell and live/dead cell discrimination, and expression of T cell-specific CD3ε expression (CD3ε^+^). Expression of γδTCR was used to identify αβ IETs (γδTCR^−^) and γδ IETs (γδTCR^+^). αβ IET populations were further discriminated by expression of CD4 and CD8α, while γδ IET populations were further discriminated by expression of CD2 and CD8α. The 8 αβ and γδ IET populations were further assessed for CD27^+/-^ expression, resulting in a total IET community composed of 16 discrete, nonoverlapping IET subpopulations. Representative images are from ileal tissue of a 4-week-old pig. **(B-D)** t-SNE dimensional reduction of gated flow cytometry data from all samples to reveal clustering of αβ and γδ IET populations **(B)**, CD27^+^ and CD27^−^ expression **(C)**, and tissue and age-specific cell frequency distributions **(D)**. In **B & C**, individual points represent single cells. Plot axes indicate t-SNE dimensions. Cells highlighted in non-grey colors coordinate with corresponding αβ and γδ IET populations **(B)** or CD27^+^ classification **(C)**. In panel **D**, areas with more cells present are indicated by red, whereas areas with less cells present are indicated by blue. Subsets of total IETs were taken from each sample (n = 990) to obtain equal cell numbers for each individual sample; an equal number of cells are present for each animal and combination of intestinal tissue and timepoint. Samples were taken from 4 animals per timepoint in each of 2 separate experiments (n = 8 per timepoint; n = 24 total).

Fluorescence intensities of cell surface markers within the total CD3ε^+^ T cell community were utilized to implement t-SNE visualization of IETs (Supplemental Fig. 4). Visualization revealed close proximities of cells assigned to each of the 8 IET populations, as defined by γδTCR, CD2, CD4, and/or CD8α expression in Fig. 2A, and variability in frequencies of cells belonging to the 8 populations (Fig. 2B). Within each of the 8 IET populations, expression of CD27 was variable based on CD27^+^ or CD27^−^ classification, but some populations were largely CD27^+^ while others were largely CD27^−^ (Fig. 2C). Thus, heterogeneity existed within IETs based on the cell surface markers assessed here. We next analyzed distribution of different IETs by observing cell distribution frequencies within the t-SNE plot across the multiple intestinal tissues and ages. Biases towards different IET populations or different CD27 expression levels within an IET population were observed between different intestinal tissues at a single timepoint or within a tissue across time, indicating both age- and intestinal location-dependent changes to IET distributions occurred (Fig. 2D). Therefore, t-SNE analysis of our data provided a large-scale impression of the overall marker-specific characteristics of IETs and demonstrated variability existed between IETs, as related to both age and intestinal site.

The overall effects of intestinal tissue, pig age, and a combination of tissue and pig age on intestinal IET community compositions were analyzed statistically by PERMANOVA, a multivariate comparisons of the frequencies of the 16 discrete IET subpopulations of each sample identified in Fig. 2A. This analysis revealed heterogeneity in the compositions of IET communities was significantly affected by tissue and age, and tissue exerted a greater influence on IET compositions than did age. A significant combinatorial influence of both tissue and age could also be observed, suggesting the effect of age was not consistent across all tissues (Fig. 3A).

**Figure 3.**
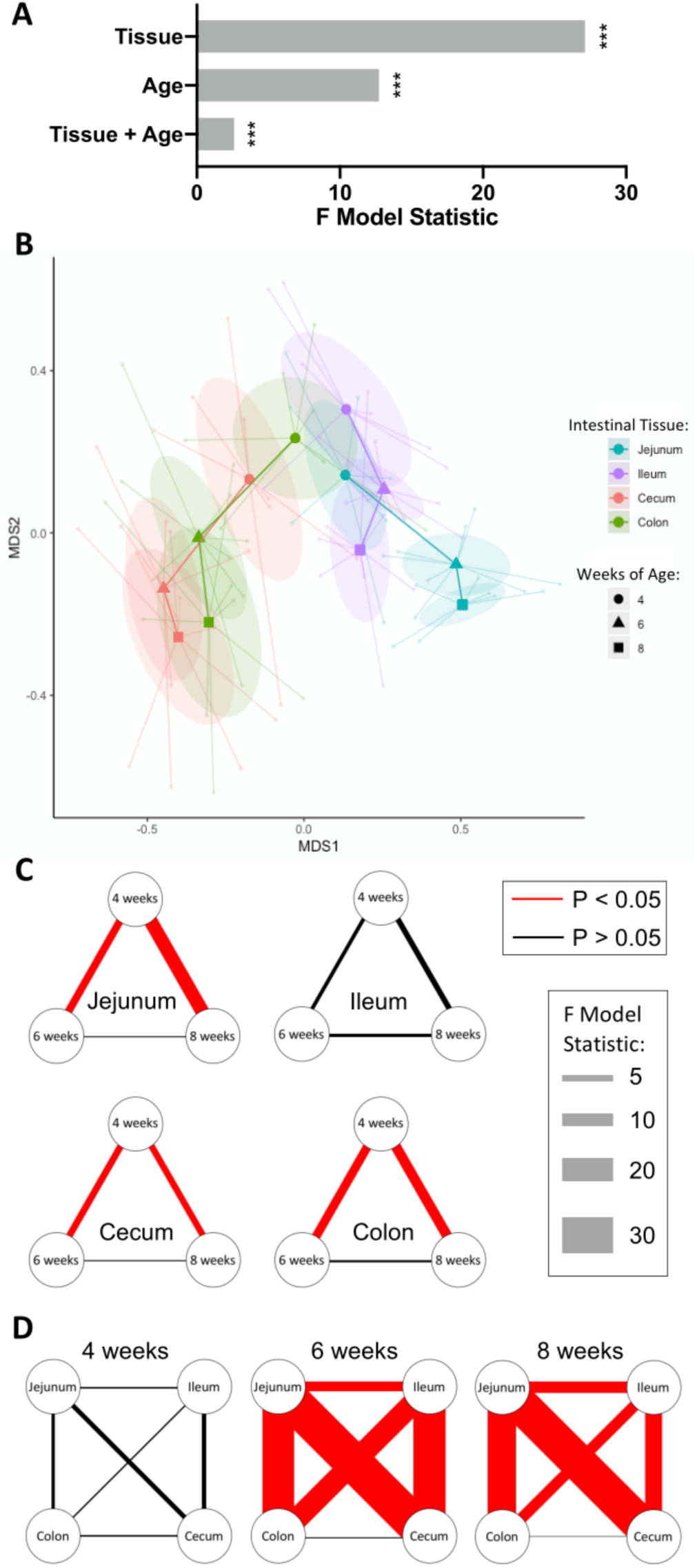
IETs diverged by intestinal location over time. **(A)** Multivariate analysis of flow cytometry data to assess tissue- and age-dependent effects on compositions of IET communities. Sample frequencies of the 16 discrete cell subpopulations defined by flow cytometry gating of αβ or γδ T cell populations and CD27 expression were analyzed for their overall tissue and age-dependent effects using PERMANOVA analysis. Greater F model statistic values represent greater magnitude of differences, while p-values < 0.05 were considered significant (* < 0.05, ** < 0.01, *** < 0.001, **** < 0.0001). **(B)** NMDS visualization of IET communities based on sample frequencies of the 16 discrete cell subpopulations defined by flow cytometry gating of αβ or γδ T cell populations and CD27 expression. Shorter distances between points represent greater similarities between samples. Centroids are plotted at the center of each 95% confidence ellipse for each sample type based on both intestinal tissue and pig age during sample collection. Centroids from each particular timepoint of a single tissue are sequentially connected chronologically. Point, ellipse, and line color is tissue-specific, while centroid shape is age-specific. **(C-D)** Visualization of pairwise PERMANOVA test results comparing IET compositions from samples between timepoints within a tissue **(C)** or between tissues within a single timepoint **(D)**. Relationships between variables are represented by connecting lines. Magnitudes of the F model statistics demonstrate the influence of a comparison and are represented by the width of the lines, while color of the line represents statistical significance (red = significant; black = not significant). P-values were corrected for multiple comparisons using the FDR method considering all tests displayed here, p-values < 0.05 were considered significant. Samples were taken from 4 animals per timepoint in each of 2 separate experiments (n = 8 per timepoint; n = 24 total).

NMDS visualization and pairwise post-hoc tests were next employed to extract results for tissue and age-specific comparisons of interest. Compositions of IET communities in jejunum, ileum, cecum, and colon were relatively similar in pigs at 4 weeks of age but diverged as age increased, with the greatest differences occurring between small and large intestinal locations (Fig. 3B). The largest changes in IET compositions within tissues over time occurred between 4 and 6 weeks of age: IET compositions in the jejunum, cecum, and colon but not ileum at 4 weeks of age were significantly different from the same sites at 6 and 8 weeks of age, while 6- and 8-week IET communities were not significantly different from each other (Fig. 3C). Divergence in IET compositions at 6 and 8 weeks of age contributed to jejunum and ileum of the small intestine becoming increasingly divergent from large intestinal tissues and, to a lesser extent, from each other (Fig. 3D). Hence, IET communities changed drastically within a single tissue mid-way through the nursery period, primarily between 4 and 6 weeks of age, and were largely defined by small or large intestinal location, although significant differences between locations within the small intestine were also detected.

### 3.3 Frequencies and phenotypes of intestinal αβ and γδ IETs differed with age and intestinal location

We next compared frequencies of different IET populations defined by FCM gating to discover the manners by which IETs differed between intestinal tissues and as pigs aged. To determine the composition of γδ versus αβ T cells within the total IET community, percentages of γδTCR^+^ cells from total CD3ε^+^ T cells obtained from our epithelial-enriched cell fractions (Supplemental Table 2) were compared across age and tissues. As no antibody reactive to the porcine αβTCR is currently available (53), CD3ε^+^γδTCR^−^ cells were inferred to be αβ T cells, while γδ T cells were identified as CD3ε^+^γδTCR^+^ (46, 54, 55). Percentages of γδTCR^+^ IETs were similar between intestinal sites at 4 weeks of age but differed between small and large intestinal tissues at both 6 and 8 weeks of age, by which times γδTCR^+^ percentages were greater in large intestinal compared to small intestinal tissues (Fig. 4A-B). Percentages of γδTCR^+^ cells decreased in jejunum between 4 and 8 weeks (p = 0.0157), whereas percentages did not change significantly in ileum, cecum, and colon over time (Fig. 4C). Thus, as animals aged, γδ T cells constituted a smaller proportion of the total IETs in small intestine compared to large intestine.

**Figure 4.**
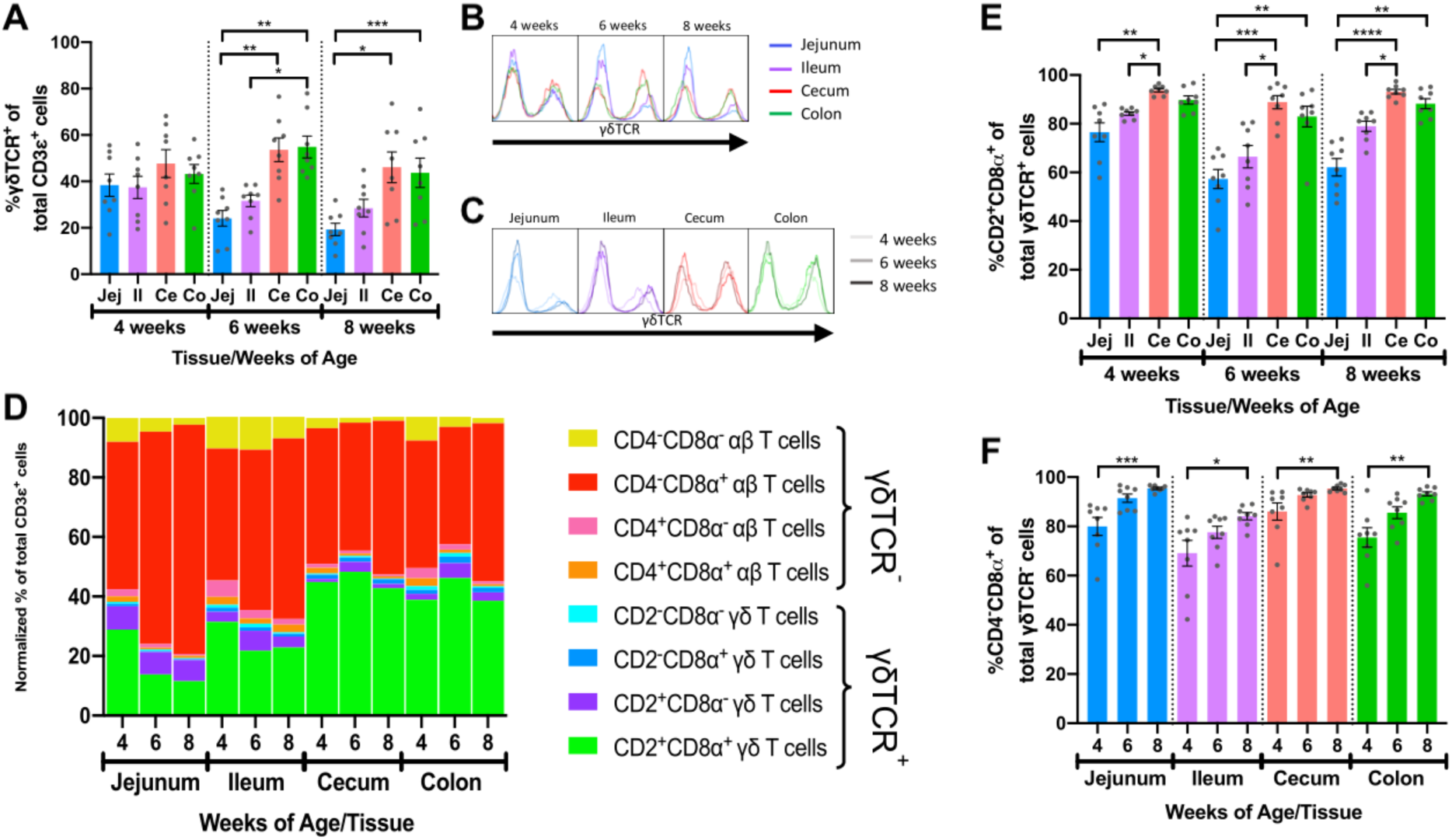
Intestinal location and age affected frequencies and phenotypes of αβ and γδ IETs. **(A)** Comparison of percentages of γδTCR^+^ cells from the total IET communities across intestinal tissues within a single timepoint. **(B-C)** Fluorescence intensities of γδTCR from the total IET communities. Frequency is proportional to unit area under the curve, and all samples are represented by equal total unit area under the curve. For fluorescence intensity curves **(B-C)**, data are acquired from merging the subsetted data of 8 samples from each age/tissue combination used for t-SNE analysis. The same data are presented in B & C to show tissue-dependent **(B)** or age-dependent **(C)** changes. **(D)** Mean normalized frequencies of αβ and γδ IET populations across intestinal tissues and timepoints. Each IET population frequency is represented by a different color, and percentages of all IET populations within a single tissue and timepoint total 100%. **(E)** Comparison of percentages of CD2^+^CD8α^+^ cells from the total γδ IET communities across intestinal tissues within a single timepoint. **(F)** Comparison of percentages of CD4^−^CD8α^+^ cells from the total αβ IET communities across timepoints within a single intestinal tissue. In **A & E**, statistical significance was determined within a single timepoint across tissues by the paired, rank-based Friedman test using all combinations of multiple comparisons within timepoint. In **F**, statistical significance was determined within a single tissue across timepoints by the rank-based Kruskal-Wallis test using all combinations of multiple comparisons within a tissue. P-values < 0.05 were considered significant (* < 0.05, ** < 0.01, *** < 0.001, **** < 0.0001). Samples were taken from 4 animals per timepoint in each of 2 separate experiments (n = 8 per timepoint; n = 24 total).

Frequencies of the γδ and αβ IET populations defined by CD2 and CD8α or CD4 and CD8α expression, respectively, varied within our total dataset (Fig. 2B). In support of this observation, the normalized frequencies of the 8 discrete γδ and αβ IET populations to the entire CD3ε^+^ IET community were calculated for samples based on intestinal tissue and age (Supplemental Table 3). Two major populations comprised a combined normalized frequency of between 78.4% to 91.7% of the total IET communities within the 12 sample groups: CD2^+^CD8α^+^ γδ T cells and CD4^−^CD8α^+^ αβ T cells (Fig. 4D).

Not only did frequencies vary between αβ and γδ IET populations, but populations of γδ or αβ IETs also appeared to vary between tissues and with age. To investigate, percentages of each of 4 IET populations within total γδ or αβ IETs (Supplemental Tables 4-5) were compared across tissues and age. γδ IET populations were defined by expression of CD2 and CD8α (Fig. 4E & Supplemental Fig. 4A-C). CD2^+^CD8α^+^ γδ T cells, which have been proposed to be a terminally differentiated cell population (29, 45, 52), comprised the largest fraction of γδ IETs in all samples, while CD2^+^CD8α^−^ γδ T cells, which are proposed to be a naïve or memory population (29, 45, 52), were the second most abundant γδ IET population (Fig. 4D). Increased percentages of CD2^+^CD8α^−^ γδ IETs within total γδ IETs (Supplemental Fig. 5A) coincided with complementary decreases in percentages of CD2^+^CD8α^+^ γδ IETs (Fig. 4E) in small intestine compared to large intestine. Moreover, differences in compositions of γδ IET populations were tissue-but not age-dependent, and, overall, CD2^+^CD8α^+^ cells were still the predominating γδ IET population within the intestinal epithelium, making up an average of between 57.3% to 93.8% of all γδ IETs, regardless of intestinal location or timepoint analyzed. Thus, the majority of γδ IETs were associated with a terminally differentiated cell phenotype (CD2^+^CD8α^+^), and γδ IETs associated with a naïve or memory phenotype (CD2^+^CD8α^−^) were more frequent in the small intestine compared to the large intestine.

Porcine αβ T cell populations were defined by CD4 and CD8α expression as presumable cytotoxic T cells (CD4^−^CD8α^+^) (47, 56), naïve helper T cells (CD4^+^CD8α^−^) (47, 48, 56), activated or memory T helper cells (CD4^+^CD8α^+^) (48, 56), and CD4^−^CD8α^−^ T cells. Porcine CD4^−^CD8α^−^ αβ T cells are not well described, but these cell are perhaps mucosal-associated invariant T (MAIT) cells (57), invariant natural killer T (iNKT) cells (58), or immature αβ T cells that will gain CD8α expression after entry into the intestinal epithelium (becoming CD4^−^CD8α^+^ αβ IETs) (7) (Fig. 4F & Supplemental Fig. 4D-F). The majority of intestinal αβ IETs were CD4^−^CD8α^+^ (average of 69.1% to 95.4%), followed in frequency by CD4^−^CD8α^−^ (1.9% to 17.3%) and then CD4^+^CD8α^+/-^ populations (0.8% to 5.0% and 0.7% to 9.5%, respectively) (Fig. 4D). As pigs aged, significant increases in CD4^−^CD8α^+^ αβ IET percentages (Fig. 4F) and concomitant decreases in all other αβ IET population percentages (CD4^+^CD8α^+^, CD4^+^CD8α^−^, and CD4^−^CD8α^−^; Supplemental Fig. 5D-F) occurred across all tissues. Findings indicate a potential increase in CD4^−^CD8α^+^ αβ IETs, decrease in other αβ IET populations (CD4^+^CD8α^+^, CD4^+^CD8α^−^, and CD4^−^CD8α^−^), or combination of both scenarios occurred at all intestinal locations as pigs aged.

### 3.4 Fewer IETs expressed CD27 in distal intestinal tract as pigs aged

CD27 is an effective phenotypic marker for functional classification of porcine T cells and is downregulated on activated/effector T cells (49-52, 59, 60). CD27 expression within intestinal IET populations was assessed to compare activation phenotypes across intestinal tissues with age (Supplemental Table 6). The 2 predominating IET populations, CD2^+^CD8α^+^ γδ IETs and CD4^−^CD8α^+^ αβ IETs, had similar patterns of CD27 expression: as pigs aged, percentages of CD27^+^ IETs decreased significantly in ileum, cecum, and colon but not jejunum (Fig. 5A-B, D-E). Decreases in percentages of CD27^+^ cells also occurred in CD2^+^CD8α^−^ γδ IETs but not any other γδ or αβ IET populations (Supplemental Fig. 6A-F). Although percentages of CD27^+^ cells decreased in ileum, decreases in percentages of CD27^+^ cells in large intestine between 4 and 6 weeks of age were more drastic. Consequently, average percentages of CD27^+^CD2^+^CD8α^+^ γδ IETs and CD27^+^CD4^−^CD8α^+^ αβ IETs were no greater than 49.0% in large intestine compared to a minimum of 64.6% in small intestine at 6 and 8 weeks of age. The data suggest the majority of IETs were not activated in 4-week-old pigs, but, as pigs continued to age, IETs of the more distal intestinal tract became activated, especially those of the large intestine.

**Figure 5.**
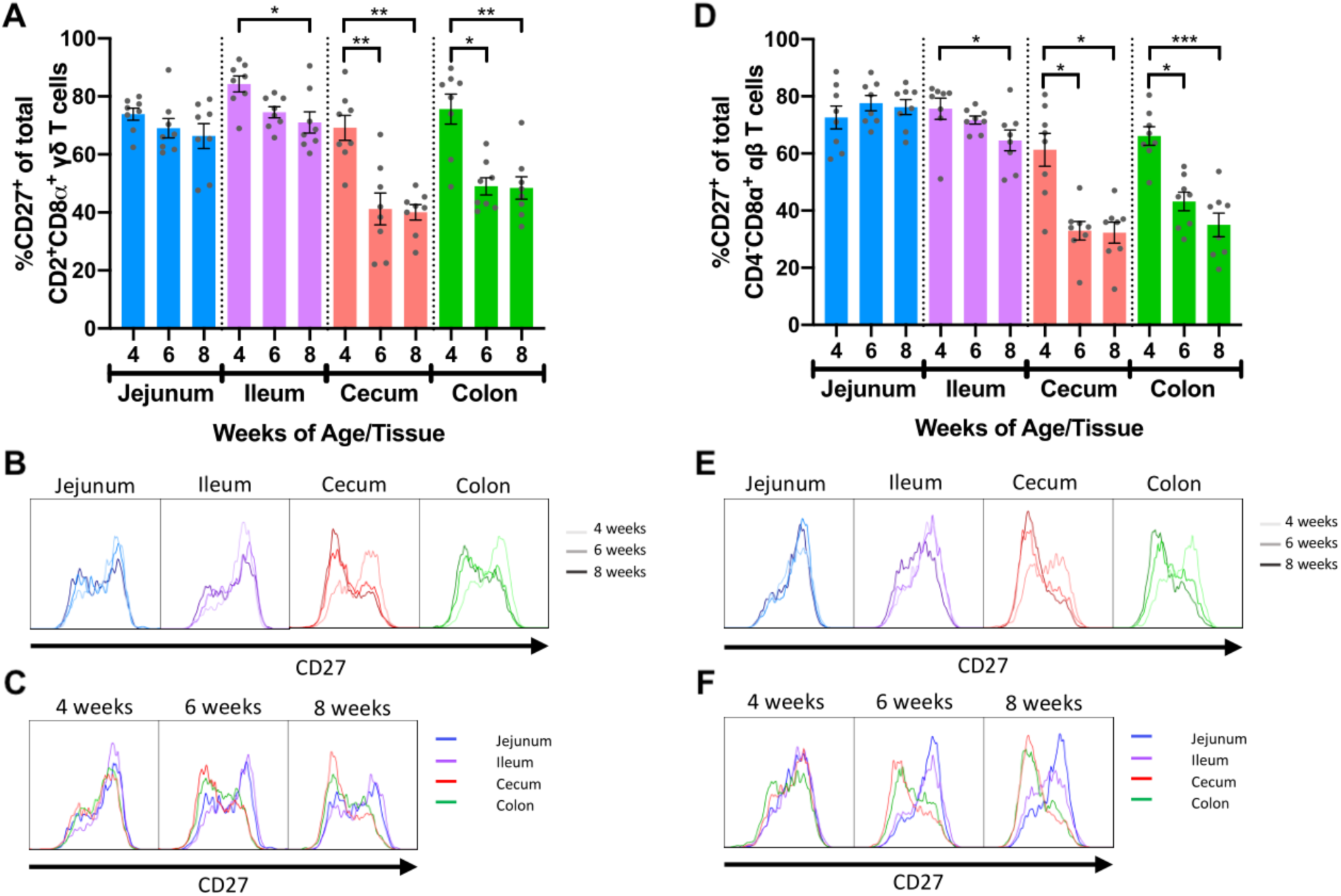
Frequencies of IETs expressing CD27 decreased with age in distal intestinal tract. **(A)** Comparisons of percentages of CD27^+^ cells within CD2^+^CD8α^+^ γδ IETs across timepoints within a single intestinal tissue. **(B-C)** Fluorescence intensities of CD27 from total CD2^+^CD8α^+^ γδ IETs within a single intestinal tissue across time **(B)** or within a single timepoint across intestinal tissues **(C). (D)** Comparisons of percentages of CD27^+^ cells within CD4^−^CD8α^+^ αβ IETs across timepoints within a single intestinal tissue. **(E-F)** Fluorescence intensities of CD27 from CD4^−^CD8α^+^ αβ IETs within a single intestinal tissue across time **(E)** or within a single timepoint across intestinal tissues **(F)**. In **B-C & D-E**, frequency is proportional to unit area under the curve, and all samples are represented by equal total unit area under the curve. For fluorescence intensity curves, data are from merging the subsetted data of 8 samples from each timepoint/tissue combination used for t-SNE analysis. The same data are presented in B & C or D & E to show age-dependent **(B, D)** or tissue-dependent **(C, E)** changes. Statistical significance was determined for percentages of total CD27^+^ cells within a single tissue across timepoints by the rank-based Kruskal-Wallis test using all combinations of multiple comparisons within a tissue. P-values < 0.05 were considered significant (* < 0.05, ** < 0.01, *** < 0.001, **** < 0.0001). Samples were taken from 4 animals per timepoint in each of 2 separate experiments (n = 8 per timepoint; n = 24 total).

## 4 Discussion

Understanding changes in IET quantities, proportional phenotypes of αβ and γδ IET populations, and expression of T cell activation marker CD27 between intestinal locations over the nursery period is important for establishing a paradigm of porcine IET maturation that may contribute to developing strategies to improve pig health and/or market performance. Herein, we demonstrate dynamics of IET maturation within the porcine intestinal epithelium arise in an age- and intestinal location-dependent manner, and our findings recapitulate and expand upon results of previous studies of porcine IETs. In line with previous work, we report IETs increased in abundance within the small intestine as pigs aged (30) and further demonstrated increased abundances of IETs within the large intestine as pigs aged. The majority of porcine IETs are CD4^−^CD8α^+^ (34), and we further show that porcine IETs belonged to both αβ and γδ T cell lineages. Overall, IETs were primarily CD2^+^CD8α^+^ γδ T cells and CD4^−^CD8α^+^ αβ T cells. Early in the nursery period (4 weeks of age), IET communities were similar throughout the intestinal tract. As pigs aged, not only did IET numbers increase, but communities became regionally specialized by 6 weeks of age, corresponding to previous findings denoting intestinal T cell communities do not resemble adult populations until between 5 to 8 weeks of age in conventional pigs (36). Moreover, shifts in IET compositions were largely due to several alterations to IET populations. First, small intestinal tissues had lower percentages of γδ IETs (and presumably greater percentages of αβ IETs) but higher proportions of CD2^+^CD8α^−^ cells (complemented by lower percentages of CD2^+^CD8α^+^ cells) within total γδ IETs than did large intestinal tissues at the later nursery stages (6 and 8 weeks of age). Second, as pigs aged, the percentages of CD4^−^CD8α^+^ αβ IETs increased in all tissues. Third, the percentages of IETs expressing CD27 decreased with age in the major IET populations (CD2^+^CD8α^+^ γδ IETs and CD4^−^CD8α^+^ αβ IETs) of the ileum and large intestine. Fourth, at later nursery stages (6 and 8 weeks of age), lower percentages of cells from the major IET populations were CD27^+^ in the large compared to small intestine. To our knowledge, this work comprises the most comprehensive and detailed analysis of porcine IETs in regards to intestinal location and nursery pig age and highlights important age- and intestinal location-associated dynamics of IET maturation to consider in future work.

Associations exist between intestinal IET phenotype data obtained from our study and previous reports of intestinal microbial diversity in pigs. The largest microbial differences in the porcine intestinal tract correlate to intestinal location, but age also correlates to microbial differences (61). Drawing parallels to the current study, intestinal location gave greater dictation to IET community composition than age, though age still had significant influence. In pigs, distinct microbial communities exist between the small and large intestine (62) but also between the jejunum and the ileum of the small intestine (63). Similarly, we detected significant differences in overall IET community compositions not only between the small and large intestine but also between jejunum and ileum. Microbial abundance and diversity increase going from proximal to distal end of the intestinal tract and as pigs age (1, 64, 65), resulting in exposure to a larger and more diverse microbially-derived antigenic repertoire in distal intestinal regions and in older pigs. Correspondingly, percentages of CD27^+^ IETs were reduced at the distal but not proximal end of the intestinal tract as pigs aged, indicating distally-located intestinal IETs may be activated at higher frequencies, and the frequency of activation accrues with age (and presumably, antigen exposure). Hence, antigenic stimulation from the microbiota should be strongly considered for potential influences on variability, maturation, and/or activation of intestinal IETs.

Ultimately, functional specialization of different intestinal segments may contribute directly or indirectly to variability in IET numbers, compositions, and activation. Variability in microbiota, dietary constituents, and immune-modulating molecules have all correlated to physiological functions at distinct intestinal locations (1). IETs were less abundant in intestinal crypts than in villi and in large compared to small intestine. In the small intestine, villi are important for increasing surface area to maximize nutrient absorption (1, 66). Hence, the small intestinal villi provide a large surface area with close contact with luminal contents, and interactions at the luminal-epithelial interface may attract greater numbers of IETs. Compared to the single mucus layer of the small intestine, the large intestine has two mucus layers, providing an additional degree of physical separation between the epithelium and luminal contents (67), and the large intestine also lacks villi. Thus, IETs located within large intestinal crypts may have less exposure to luminal contents, resulting in fewer IETs present there. Microbes in the large intestine are also important bioreactors that ferment indigestible components into metabolites that serve as fuel or signaling molecules for the host. To maintain beneficial microbes, a symbiotic relationship is established, and microbes are tolerated yet tightly regulated by the host (68, 69), perhaps through IETs in close proximity with these microbes being primed for activation. It’s unclear why lower percentages of IETs expressed CD27 in large intestine compared to small intestine as pigs aged, indicating greater percentages of IETs were activated in the large intestine compared to the small intestine. However, differing compositions and abundances of microbial species present in the large intestine compared to the small intestine may play a role in IET activation. Moreover, large intestinal IETs may be exposed to a different repertoire of soluble factors than IETs of the small intestine, such as microbially-derived components including lipopolysaccharide (LPS) or microbially-constructed metabolite products including short-chain fatty acids (SCFAs).

In peripheral blood and non-intestinal tissues, the presence or absence of CD8α expression within porcine CD2^+^ γδ T cells is associated with different functional states. CD2^+^CD8α^+^ γδ T cells exhibit greater expression of the effector T cell transcription factor T-bet (52), greater cytokine production (45), and expression patterns of cell surface molecules indicative of T cell activation (29, 49) when compared to CD2^+^CD8α^−^ γδ T cells. Resultingly, CD2^+^CD8α^−^ γδ T cells are a proposed naïve or memory cell population, while CD2^+^CD8α^+^ γδ T cells are proposed to be a terminally differentiated population (45, 52). Moreover, porcine CD2^+^CD8α^−^ γδ T cells can gain expression of CD8α following *in vitro* IL-2 stimulation (46). In humans and rodents, IETs may gain surface expression of the CD8αα homodimer upon activation (7), supporting the notion that porcine CD2^+^CD8α^+^ γδ T cells are a terminally differentiated cell population arising from CD2^+^CD8α^−^ γδ T cells. In our study, greater percentages of γδ IETs expressed CD8α in the distal compared to proximal intestinal tract. Expression of the CD8αα homodimer on T cells is implicated in increasing the threshold for T cell receptor-mediated activation (70-72). Hence, CD8α expression in our proposed terminally differentiated CD2^+^CD8α^+^ γδ IETs may indicate an increased threshold for TCR-mediated activation, suggesting a more regulatory or tolerant phenotype for CD2^+^CD8α^+^ γδ IETs compared to CD2^+^CD8α^−^ counterparts. Whether similar phenomena occur in pigs is unknown but supported by previous *in vitro* and *in vivo* work in gnotobiotic pigs (23, 73). Wen et al. demonstrate ileal CD2^+^CD8α^−^ γδ IETs secreted higher levels of IFN-γ, secreted lower levels of IL-10, and expressed lower levels of regulatory transcription factor Foxp3 compared to CD2^+^CD8α^+^ γδ IETs (73). Moreover, frequencies of CD2^+^CD8α^+^ γδ T cells increased in ileum of gnotobiotic pigs following colonization with probiotic *Lactobacilli* or infection with human rotavirus, while CD2^+^CD8α^−^ frequencies decreased (23). Therefore, it is plausible that γδ IETs in the more distal intestinal tract have an increased threshold for TCR-mediated activation, giving way to a more tolerogenic profile. The same might also hold true for αβ IETs; however, whether CD4^−^CD8α^+^ αβ IETs expressed CD8αα, CD8αβ, or a combination of both cannot be determined from our data.

In the United States, pigs are typically weaned at ∼21 days of age and immediately proceed to the nursery period (3 to 10 weeks of age) of production thereafter. The weaning period is considered to be one of the most stressful life events for pigs. Weaning involves cessation of passive immune transfer of milk-derived immunoglobulins by abrupt removal of piglets from the sow; introduction of new social and environmental stressors from inter-litter mixing and transport to a new facility; and introduction to a solid-food diet (38, 39). In addition, animals are exposed to a plethora of new environmental, microbial, and dietary antigens that may induce age-associated changes to intestinal IETs. In humans and rodents, major naturally-occurring IET populations include CD4^−^CD8α^+^CD8β^−^ αβ T cells (CD8αα^+^ αβ IETs), which express the CD8αα homodimer rather than the CD8αβ heterodimer, and γδ T cells (2, 7). Natural IETs are recruited to the epithelium in an antigen-independent manner and do not show increased recruitment with antigen exposure or age (2). Induced IETs, on the other hand, are predominately CD4^−^CD8α^+^CD8β^+^ αβ T cells (CD8αβ^+^ αβ IETs) expressing the CD8αβ coreceptor (2, 7). Induced IETs encounter their cognate antigen, then are recruited to the intestinal epithelium and reside within the epithelial layer as antigen-experienced effector or memory cells (8, 74). Induced IETs increase with age in association with increased antigen exposure (2). Hence, increased IET numbers observed in our study may be related to the recruitment of induced IETs associated with exposure to increased antigenic load and/or diversity as pigs aged. Increases in induced IET recruitment as antigen is experienced could account for overall increases in IET numbers observed as pigs aged, as well as increased percentages of CD4^−^CD8α^+^ αβ IETs observed across time at all intestinal sites. Moreover, decreased percentages of γδ IETs in the small intestine compared to large intestine may be attributed to greater recruitment and abundance of induced CD8αβ^+^ αβ IETs in the small intestine. To our knowledge, natural and induced IETs have not been characterized in the pig intestinal tract, but knowledge of CD8β expression for αβ T cells required to further investigate presumable induced or natural IET phenotypes based on human and rodent data is lacking from our data.

## 5 Conclusions

Overall, we demonstrated heterogeneity in IET numbers, compositions, and activation phenotypes between small and large intestinal tissues and across age in nursery pigs. IET communities were largely similar between intestinal sites early in the nursery phase; however, tissue-specific divergence occurred as pigs aged, indicating the nursery period is a critical time of intestinal IET maturation in conventional pigs. Divergence in IET communities was evident by variation in cell numbers, γδ versus αβ T cell compositions, frequencies of γδ and αβ IET populations, and CD27 expression. Due to the uniqueness of IETs by intestinal location and pig age, results pertaining to IETs should not be generalized, but rather the variables of age and intestinal location should be strongly considered. Our findings are based on cellular phenotypes, and functional significance remains to be shown. In this regard, caution should be taken when applying functional characteristics based from research performed using T cells that are not of intestinal epithelial origin. Though not addressed by our study, additional factors such as intestinal microbiota, weaning age, antibiotic usage, environmental stress, diet, animal market performance, and disease, should be strongly considered for their correlations with or impacts on the age- and intestinal location-dependent IET communities defined herein.

## 6 Ethics Statement

All procedures adhered to the ethical and humane use of animals for research and were approved by the Iowa State University Institutional Animal Care and Use Committee (IACUC #18-351).

## 7 Author Contributions

JEW, NKG, and CLL designed experiments. JEW, ZFB, NKG, and CLL collected samples. JEW, ZFB, KAB, and CLL performed experiments. JEW and JMT analyzed data. JEW wrote the manuscript. All authors reviewed and edited the manuscript.

## 8 Funding

This research was supported by (1) appropriated funds from USDA-ARS CRIS project 5030-31320-004-00D and (2) an appointment to the Agricultural Research Service (ARS) Research Participation Program administered by the Oak Ridge Institute for Science and Education (ORISE) through an interagency agreement between the U.S. Department of Energy (DOE) and the U.S. Department of Agriculture (USDA). ORISE is managed by ORAU under DOE contract number DE-SC0014664. All opinions expressed in this paper are the authors’ and do not necessarily reflect the policies and views of USDA, ARS, DOE, or ORAU/ORISE.

## 9 Conflict of Interest Statement

The authors declare that the research was conducted in the absence of any commercial or financial relationships that could be construed as a potential conflict of interest.

## 10 Acknowledgements

We thank the following for their outstanding contributions to this work: Trey Faaborg and the Iowa State University Swine Nutrition Farm for animal care; Jamison Slate, Jessica Jasper, Emma Helm, and Carson De Mille for necropsy assistance; Sam Humphrey for flow cytometry expertise; Judith Stasko and Adrienne Shircliff for histology services; and Michael Marti for creation of Figure 1A.

## Notes

https://github.com/jwiarda/Intraepithelial_T_cells

