## Supplementary Figures for "Intraepithelial T cells diverge by intestinal location as pigs age"

### *Supplementary Material*

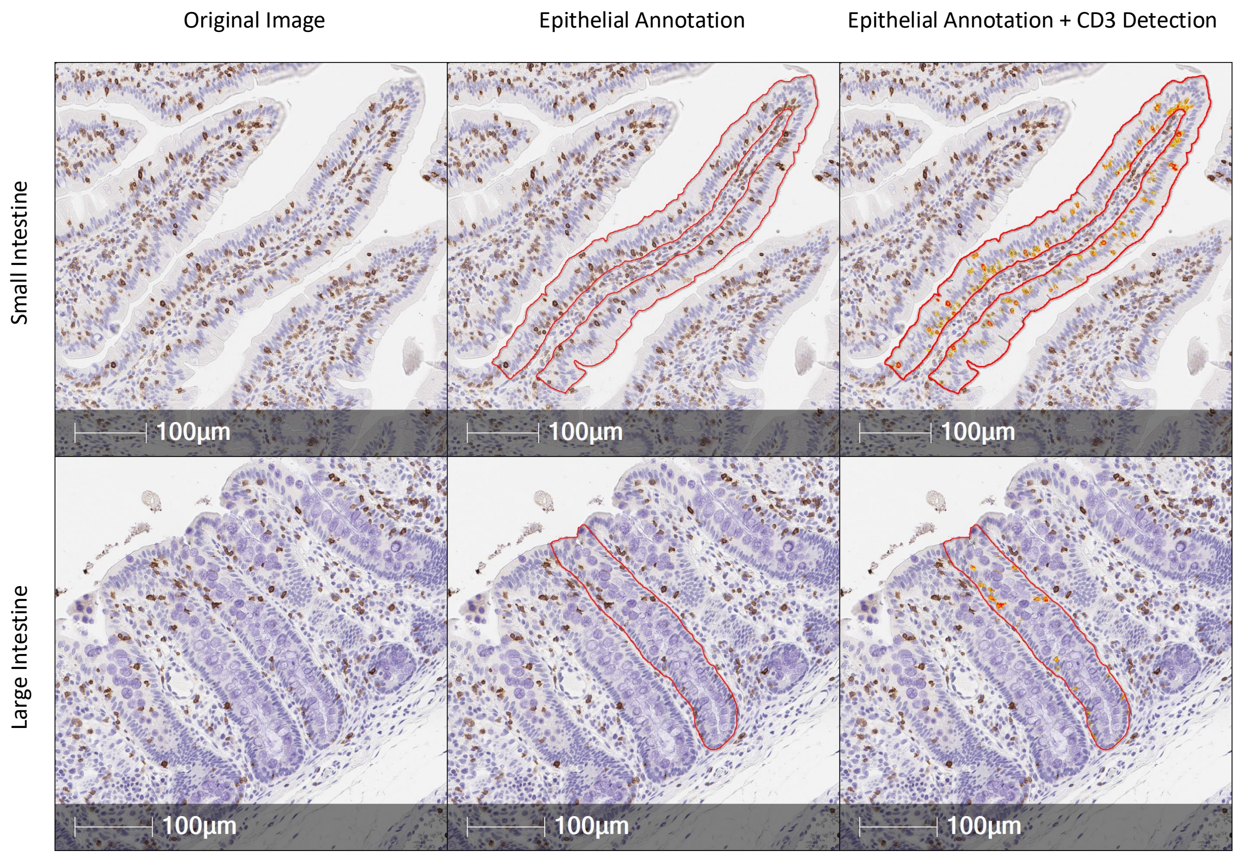


**Supplementary Figure 1. Representative images of quantitative tissue immunohistochemistry staining analysis.** From original tissue images (left), small intestinal villi (top) and large intestinal crypt (bottom) epithelia were annotated (middle) and then analyzed to detect CD3 staining (right). Surface area stained positive for CD3 (yellow, orange, and red areas detected in right panels) was divided by total surface area within the annotation layer (middle panel) to calculate surface area staining positive for CD3 within epithelial annotation layers. A total of 3 intestinal villi or crypts were analyzed from each intestinal tissue of each animal, and surface areas from the 3 areas were averaged together for an overall quantification of each tissue. Samples were taken from 4 animals per timepoint in each of 2 separate experiments (n = 8 per timepoint; n = 24 total).

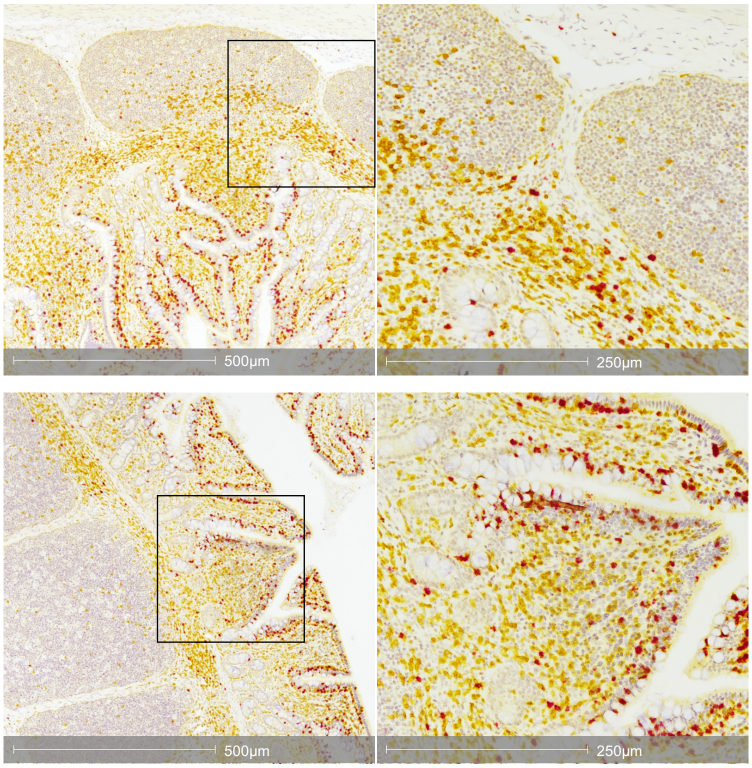


**Supplementary Figure 2. Distribution of γδ and αβ T cells in submucosa, lamina propria, and epithelium.** Image of T-cell specific CD3 protein staining (yellow) and γδ T cell-specific *TRDC* transcript staining (red) in pig ileum at 8 weeks of age. *TRDC*$^{+}$ and *TRDC*$^{-}$CD3$^{+}$ cells were both present in Peyer's patch (top) and dome (bottom) regions.

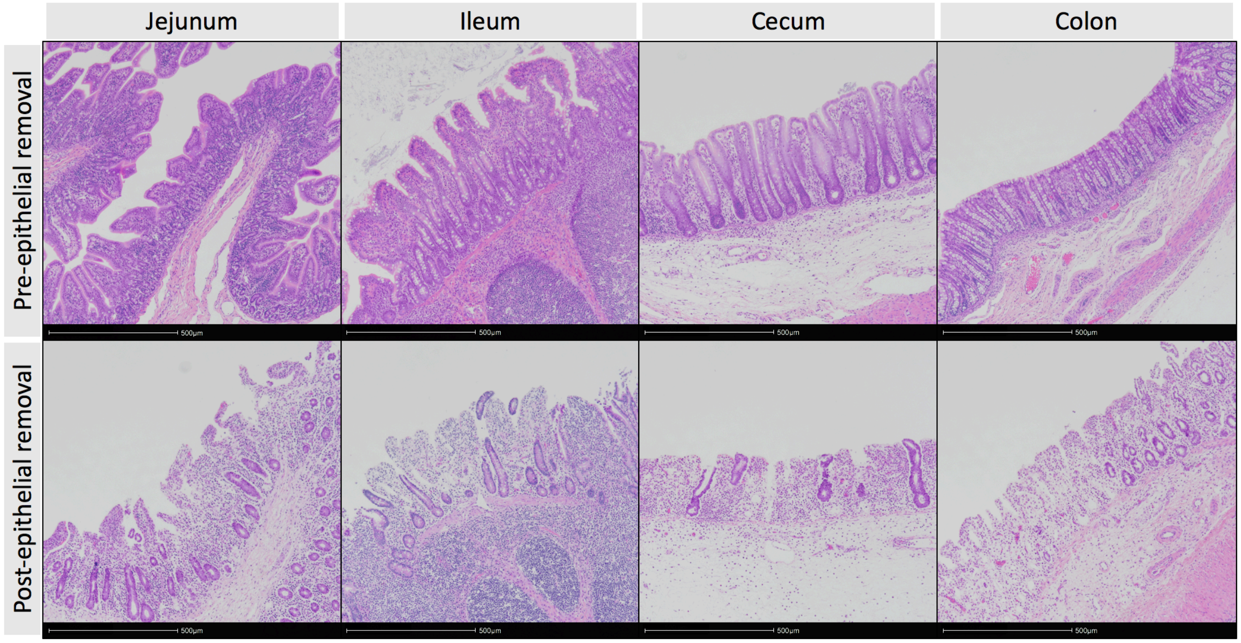


**Supplementary Figure 3. Incubation for epithelial removal efficiently removed epithelial cells while leaving lamina propria and submucosa intact in pig intestinal tissues.** Representative images of intestinal tissues pre- (top) and post-incubation (bottom) in epithelial removal solution from 6-week-old pig.

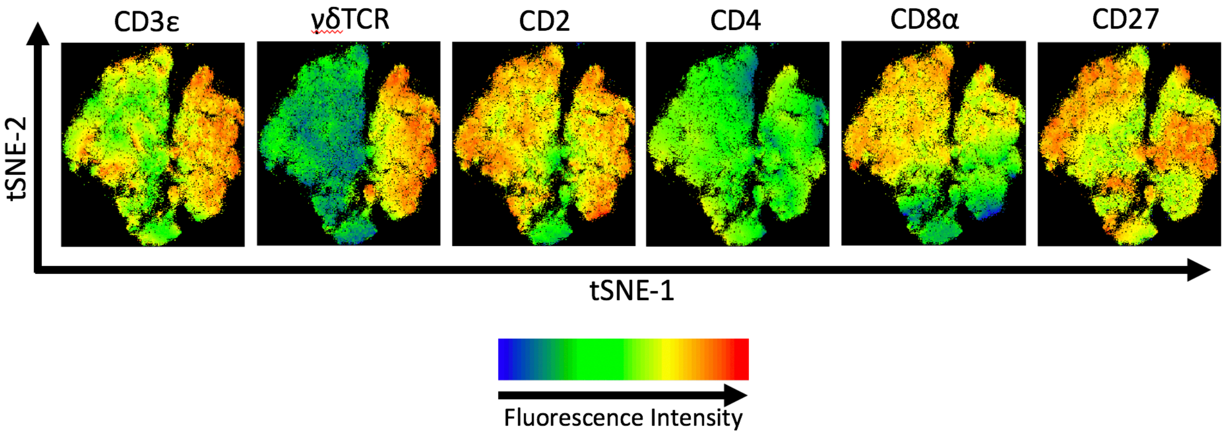


**Supplementary Figure 4. Fluorescence intensities of flow cytometry markers allow t-SNE clustering of IETs.** Fluorescence intensities of CD3ε, γδTCR, CD2, CD4, CD8α, and CD27 across equal numbers of CD3$\varepsilon^{+}$ cells from each sample of jejunum, ileum, cecum, and colon from 4-, 6-, and 8-week-old pigs. Fluorescence intensities of these markers were used to cluster cells using t-SNE. Samples were taken from 4 animals per timepoint in each of 2 separate experiments (n = 8 per timepoint; n = 24 total).

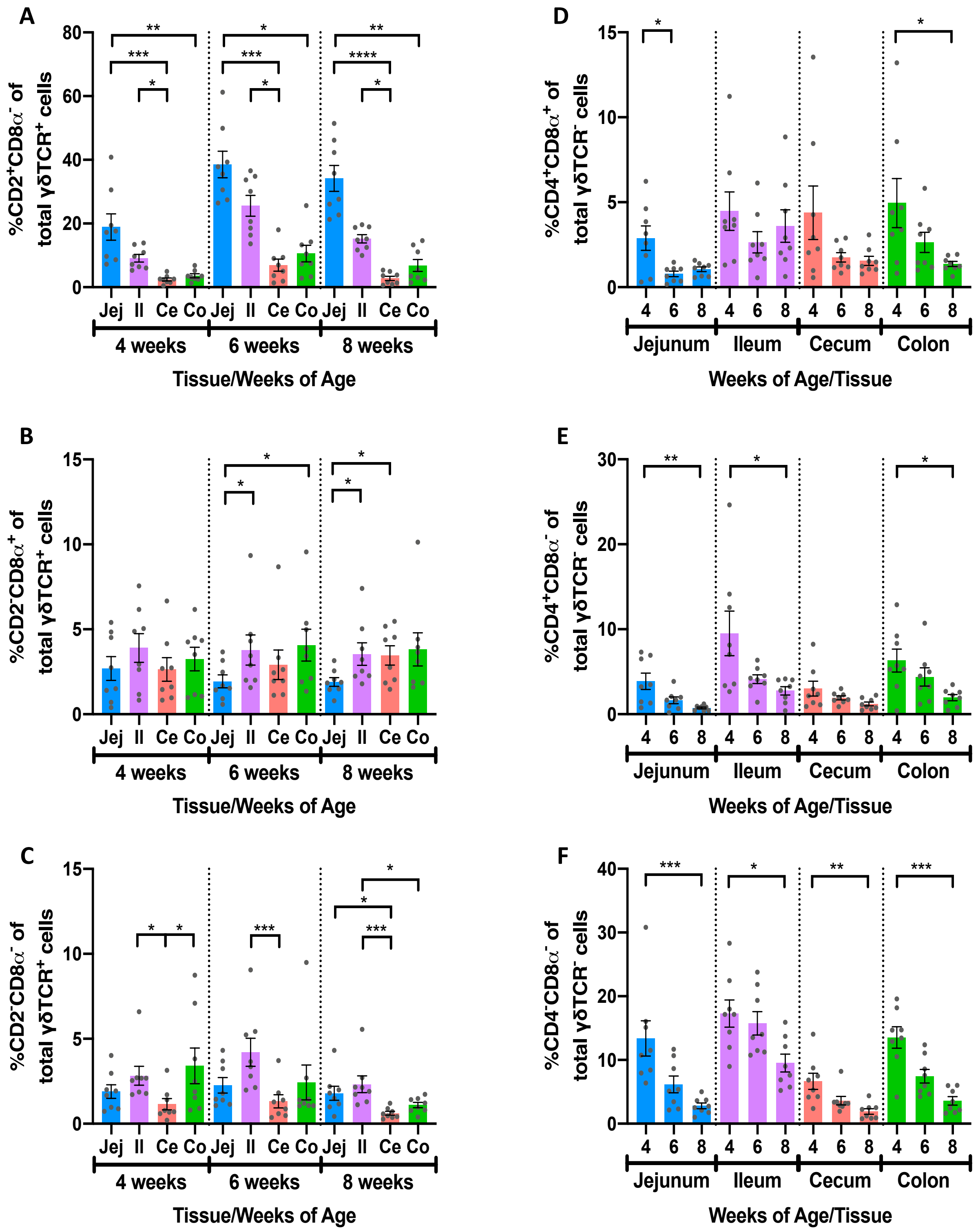


**Supplementary Figure 5. IET population frequencies within total γδ or αβ IETs. (A-C)** Comparison of percentages of $CD2^{+}CD8\alpha^{-}$ **(A)**, $CD2^{-}CD8\alpha^{+}$ **(B)**, and $CD2^{-}CD8\alpha^{-}$ **(C)** IET populations from total γδ IETs across intestinal tissues within a single timepoint. Statistical significance was determined within a single timepoint across tissues by the paired, rank-based

**Supplementary Figure 5 (continued)** Friedman test using all combinations of multiple comparisons within a timepoint. **(D-F)** Comparison of percentages of $CD4^{+}CD8\alpha^{+}$ **(D)**, $CD4^{+}CD8\alpha^{-}$ **(E)**, and $CD4^{-}CD8\alpha^{-}$ **(F)** IET populations from total αβ IETs across timepoints within a single intestinal tissue. Statistical significance was determined within a single tissue across timepoints by the rank-based Kruskal-Wallis test using all combinations of multiple comparisons within a tissue. P-values < 0.05 were considered significant (* < 0.05, ** < 0.01, *** < 0.001, **** < 0.0001). Samples were taken from 4 animals per timepoint in each of 2 separate experiments (n = 8 per timepoint; n = 24 total).

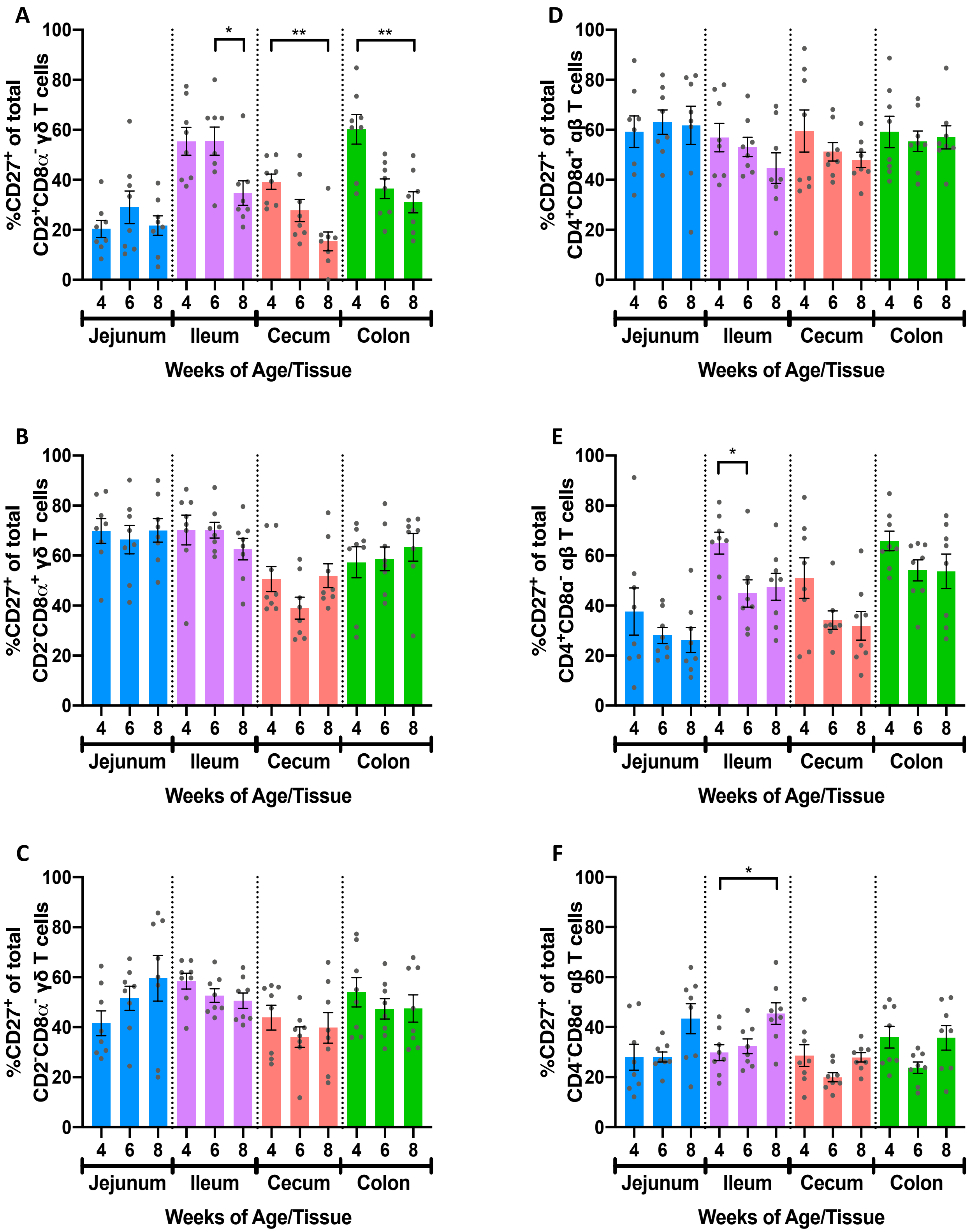


**Supplementary Figure 6. Frequencies of CD27$^{+}$ cells within γδ and αβ IET populations.** Comparison of percentages of CD27$^{+}$ cells from the total populations of CD2$^{+}$CD8α$^{-}$ **(A)**, CD2$^{-}$

**Supplementary Figure 6 (continued)** CD8α$^{+}$ **(B)**, and CD2$^{-}$CD8α$^{-}$ **(C)** γδ IETs or CD4$^{+}$CD8α$^{+}$ **(D)**, CD4$^{+}$CD8α$^{-}$ **(E)**, and CD4$^{-}$CD8α$^{-}$ **(F)** αβ IETs. Statistical significance was determined within a single tissue across timepoints by the rank-based Kruskal-Wallis test using all combinations of multiple comparisons within a tissue. P-values < 0.05 were considered significant (* < 0.05, ** < 0.01, *** < 0.001, **** < 0.0001). Samples were taken from 4 animals per timepoint in each of 2 separate experiments (n = 8 per timepoint; n = 24 total).
